# Single-molecule nanophotonic resolution of binding dynamics from apo to fully liganded for a cyclic nucleotide-gated ion channel in cell-derived vesicles

**DOI:** 10.64898/2026.05.24.727213

**Authors:** Tapas Haldar, Draegan Watson, Cecilia M. Borghese, Zaid Ahmed, Perla A. Peña Palomino, Susanne Ressl, Audrey C. Brumback, Marcel P. Goldschen-Ohm

## Abstract

Many biological macromolecules are activated upon ligand binding at multiple specific binding domains. However, how these domains interact and the transient intermediate conformations that connect binding events to protein activity are typically unknown. Ensemble-averaged measures over stochastic binding events are challenged to resolve the underlying asynchronous dynamics. Single-molecule resolution of these dynamics offers an attractive approach to investigate the ligand-activation process. Optical methods using fluorescently labeled ligands enable observation of individual binding events that report on the energetics of early ligand-bound conformational changes. However, diffraction-limited microscopy limits these methods to low ligand concentrations, often below what is required for physiologically relevant activation. Here, we overcome this limitation using nanophotonic zero-mode waveguides to observe the sequential binding of a fluorescent cyclic nucleotide to each of four subunits in TAX-4 cyclic nucleotide-gated ion channels in cell-derived vesicles. Our observations suggest that binding at one domain positively promotes binding at other domains, and that binding induces an isomerization of the binding domain in individual subunits which we attribute to a sequence of pre-activated intermediate states. This approach provides a broadly applicable tool to dissect the energetic landscape of ligand-binding at macromolecules in native cell membranes.

## Introduction

Molecular association through specific recognition events lies at the heart of biological signaling, orchestrating the precise communication required for life’s complex processes. Such interactions, whether between proteins, nucleic acids, or small molecules, initiate cascades that regulate cellular function, development, and responses to environmental cues. Despite their fundamental importance, the underlying dynamics and energetics of individual ligand binding events that precede and drive downstream protein activity remain largely elusive, in part because conventional biochemical and biophysical approaches typically infer these molecular processes indirectly.

Traditional measures of protein activity—such as ion flux, conformational change, or enzymatic turnover—report only the end results of binding, without revealing the sequence and nature of individual association events. Similarly, ensemble-averaged binding assays, while powerful for characterizing global properties, obscure asynchronous and heterogeneous dynamics by averaging over populations of molecules, effectively obscuring short-lived, partially liganded intermediates and distinct per molecule activation sequences. Consequently, such bulk approaches are often insufficient to uniquely resolve the stochastic trajectories and rare intermediate states that connect ligand binding to protein activation (Hines, Middendorf, and Aldrich 2014).

Single-molecule fluorescence methods have emerged as transformative tools to overcome these limitations, enabling direct visualization of individual binding events and conformational transitions in real time (Goldschen-Ohm et al. 2016; Patel et al. 2021; White et al. 2021; Schmauder et al. 2011; Idikuda et al. 2025; Chowdhury, Goldsmith, and Chanda 2026). These approaches are especially relevant for dissecting the activation mechanisms of membrane proteins such as cyclic nucleotide-gated (CNG) ion channels, which play pivotal roles in sensory transduction, neuronal excitability, and signal integration across diverse cell types (J. Zheng and Trudeau 2015; Komatsu et al. 1996; Coburn et al. 1998). CNG channels, found in both vertebrate and invertebrate systems, transduce chemical signals into electrical responses by gating in response to binding cyclic nucleotides at four intracellular binding domains—one per subunit in the tetrameric channel (X. Zheng et al. 2020; Taraska and Zagotta 2007; Young, Sciubba, and Siegelbaum 2001).

We previously developed an approach using micro-mirror total internal reflection fluorescence (TIRF) microscopy to resolve individual binding events of a fluorescent cGMP analog 8-DY547-cGMP (fcGMP) (Biskup et al. 2007) at single TAX-4 CNG channels immobilized in cell-derived vesicles (Patel et al. 2021). This work revealed key insights into early binding steps and partial activation, yet was limited to detecting only the first two of four binding events due to background fluorescence at higher ligand concentrations—a constraint imposed by conventional optical methods and their relatively large excitation volumes (White et al. 2023).

In the present study, we address these challenges by integrating nanophotonic zero-mode waveguides (ZMWs) with purified and extruded cell-derived vesicles containing TAX-4 CNG channels. ZMWs, utilized largely for DNA sequencing and protein-DNA interactions, confine the optical excitation volume to the zeptoliter scale, dramatically suppressing background from freely diffusing ligand and enabling single-molecule detection at physiologically relevant, micromolar ligand concentrations (Levene et al. 2003; Eid et al. 2009; Uemura et al. 2010; Goldschen-Ohm et al. 2016; White et al. 2021; Goldschen-Ohm et al. 2017). Cell-derived vesicles eliminate solubilization of the membrane protein in detergent or other synthetic environments that can affect protein function (Fox-Loe, Moonschi, and Richards 2017; Moonschi et al. 2018). As a result, we achieve sequential, real-time observation of fcGMP binding at all four channel subunits, providing unprecedented resolution of cooperative binding dynamics for a CNG channel in a native lipid context.

Our observations show that sequential binding to TAX-4 channels is a positively cooperative process. Kinetic modeling further suggest that binding induces an isomerization of the binding domain which we attribute to a pre-active intermediate state. These advances establish a broadly applicable framework for dissecting the energetic landscape of ligand association in membrane protein complexes and illuminate fundamental principles of cooperative activation in a CNG channel.

## Results

### Immobilization of TAX-4 CNG channels in cell-derived vesicles within ZMW nanophotonic arrays

We previously investigated binding of a fluorescent cGMP analog (fcGMP) (Biskup et al. 2007) to individual TAX-4 CNG channels in cell-derived vesicles using TIRF excitation of vesicles immobilized on a passivated glass coverslip (Patel et al. 2021). In this approach, TIRF limits the optical excitation volume to within a few hundred nanometers of the surface, thereby reducing the background fluorescence from freely diffusing fcGMP in the bulk solution to a level enabling resolution of single fcGMP molecules at up to low nanomolar concentrations of fcGMP (Goldschen-Ohm et al. 2017; White et al. 2023). Although this enabled observation of mono-and di-liganded bound conformations, reliable observation of tri-and quad-bound conformations could not be well resolved with TIRF as the increased background fluorescence at the higher nanomolar to micromolar fcGMP concentrations required challenged single fcGMP resolution.

To overcome this limitation, we used ZMWs which have previously enabled resolution of binding at up to micromolar concentrations in related HCN channels (Goldschen-Ohm et al. 2016, 2017; White et al. 2021; Idikuda et al. 2025; White et al. 2023). Briefly, ZMWs are nano-sized apertures in a thin metal film on an optical surface such as a glass coverslip. Light having a wavelength more than double the aperture diameter does not propagate through the aperture but can still excite fluorophores inside the aperture within a few tens of nanometers of the optical surface. This effectively reduces the optical excitation volume to the zeptoliter (10^-21^ liter) scale, much smaller than for TIRF, thereby enabling single fluorophore resolution near the bottom of the aperture with up to micromolar concentrations of fluorescent species in the bulk solution (Levene et al. 2003).

In order to avoid solubilization in non-native detergents, we generated plasma membrane vesicles from cells expressing TAX-4 channels with eGFP fused to their N-terminus for optical localization (eGFP–TAX-4) as previously described (Patel et al. 2021). However, the relatively broad size distribution of these vesicles from a few tens of nanometers up to several microns in diameter is largely incompatible with entry into ZMW apertures having 100-150 nm diameters. To limit the vesicle size distribution, we sequentially extruded vesicles through 50 nm followed by 30 nm pores resulting in a large fraction of vesicles having diameters of 50-100 nm (**Fig. S1**). We then attempted to immobilize these vesicles at the bottom of large arrays of 100-150 nm diameter ZMW apertures functionalized with anti-GFP nanobodies in the same way as previously described for chambers without ZMWs (Patel et al. 2021). However, only a very small fraction of ZMW apertures exhibited GFP fluorescence indicative of successful loading of a vesicle containing eGFP–TAX-4 oriented with its binding domains accessible to the bath (**Fig. S2**).

Given that the vesicle sample includes vesicles with or without eGFP–TAX-4 as well as vesicles oriented either inside-or outside-out, we hypothesized that apertures may be getting occluded by either empty vesicles (i.e., without TAX-4) or vesicles where TAX-4 is oriented with its binding domain trapped within the vesicle. To address this, we generated a new construct to purify TAX-4-containing vesicles having exposed binding domains. This TAX-4 construct has an ALFA tag followed by the fluorescent protein mStayGold (mSG) fused to the N-terminus, and a PreScission (PS) cleavage site followed by a Twin-Strep® (TS) tag fused to the C-terminus (ALFA–mSG–TAX-4–PS–TS; **Fig. 1**). Note that both N-and C-termini are intracellular, and that the cyclic nucleotide binding domain (CNBD) is also in the C-terminal domain. We opted for mSG due to its brighter and longer-lived fluorescence as compared to eGFP (Ivorra-Molla et al. 2024; Ando et al. 2024), which necessitated another tag for immobilization (i.e., ALFA). The PS and TS motifs are used for purification.

**Figure 1.**
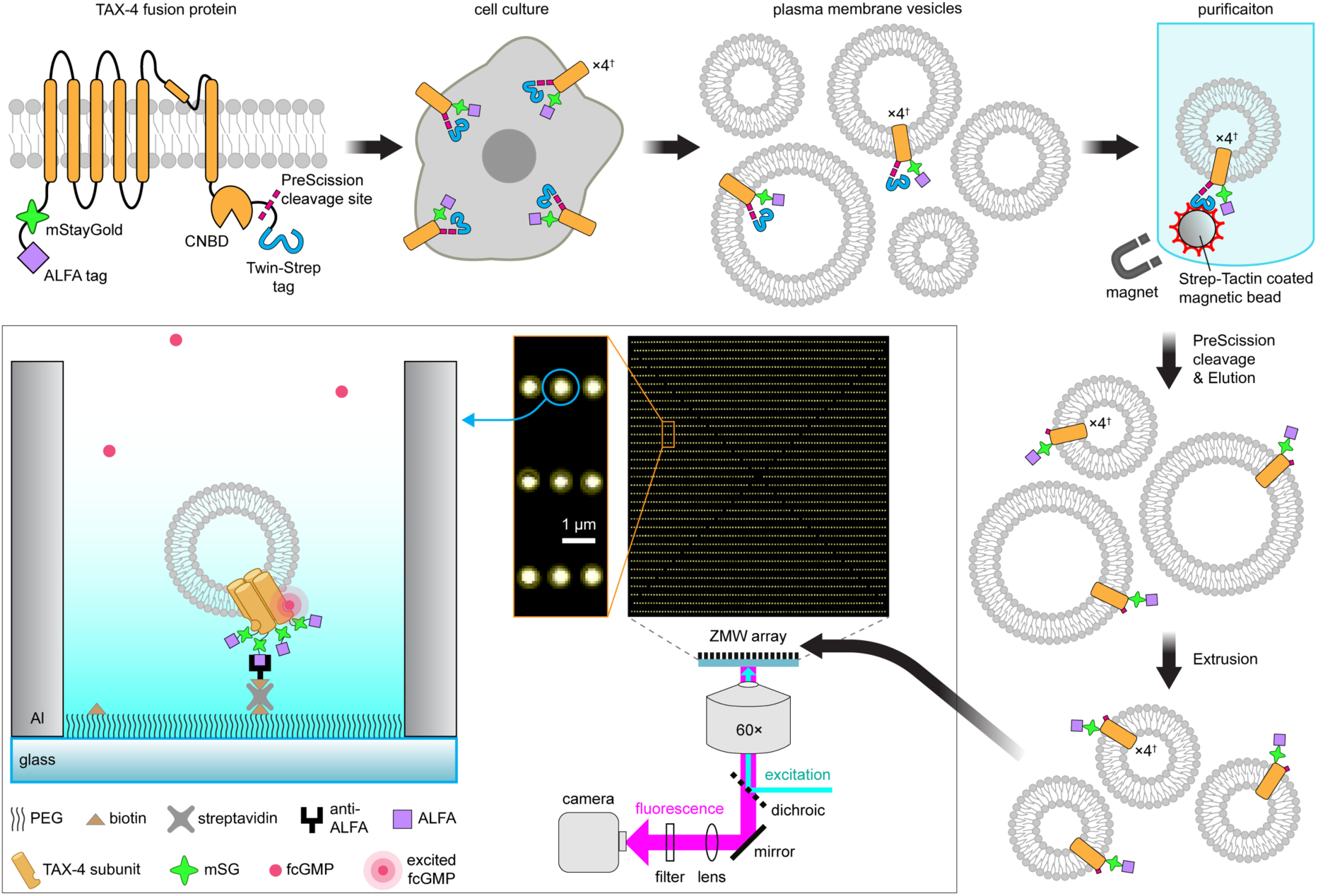
Imaging TAX-4 ion channels in cell-derived vesicles within ZMWs. Schematic of sample preparation and imaging workflow. From upper left to lower right: Topology for the TAX-4 subunit fusion protein expressed in HEK-293T cells. Plasma membrane vesicles are derived from cells using nitrogen cavitation and gradient ultracentrifugation as described previously (Patel et al. 2021; Fox-Loe, Moonschi, and Richards 2017; Moonschi et al. 2018). Vesicles containing TAX-4 channels with their N-and C-termini oriented outwards to the bulk solution were affinity purified using Strep-Tactin magnetic beads to bind the Twin-Strep tag and eluted following cleavage of the Twin-Strep tag at an upstream PreScission site. Purified vesicles were sequentially extruded through 50 and 30 nm pores to obtain vesicles with diameters ∼50-100 nm. Extruded vesicles were loaded onto ZMW arrays (middle) and immobilized at the bottom of 100-150 nm diameter ZMW apertures by anti-ALFA nanobodies tethered to a PEG layer via biotin-streptavidin interactions (bottom left). Fluorescence from the entire array of ∼3000 ZMW apertures (brightfield image shown) was recorded simultaneously under epi illumination on a sCMOS camera. Note that cartoon illustrations of TAX-4, vesicles, magnetic beads, and ZMW aperture are not to scale. ^†^To simplify visualization, only a single subunit of the tetrameric channel is shown in vesicle preparation schematics.

The ALFA–mSG–TAX-4–PS–TS fusion construct expressed in HEK-293T cells with many cells showing fluorescence indicative of expression in the plasma membrane (**Fig. S3**). Plasma membrane vesicles prepared from HEK-293T cells expressing ALFA–mSG–TAX-4–PS–TS were affinity purified using Strep-Tactin® magnetic microbeads which specifically bind the TS tag (**Fig. 1**). Briefly, vesicles were incubated with the magnetic beads to bind vesicles containing ALFA–mSG–TAX-4–PS–TS with the TS tag oriented outward. Vesicles lacking the fusion protein, or those with the TS tag trapped within the vesicle, should not bind and were removed by washing. Subsequently, we eluted bound vesicles by cleavage at the PS site using PreScission protease. The resulting eluate should contain only those vesicles with ALFA–mSG–TAX-4 having both the ALFA tag and CNBDs accessible to the bath. The efficiency of the purification was approximately 30% as assessed by quantifying mSG fluorescence from samples taken at various stages of the purification (**Fig. S4**). Purified ALFA–mSG–TAX-4 expressing vesicles were then extruded as described above and immobilized in the bottom of ZMW apertures functionalized with anti-ALFA nanobodies (**Fig. 1**). This approach enabled much more efficient loading of ZMW apertures as evidenced by the increased fraction of apertures exhibiting mSG fluorescence (**Fig. S2**). Furthermore, this immobilization was specific as almost no loading was observed for ZMWs lacking anti-ALFA nanobodies (**Fig. S5**).

### Resolving the dynamics of sequential binding to all four subunits at individual TAX-4 CNG channels

For optical detection of ligand binding events, the sample chamber was bathed with a fluorescent cGMP analog 8-DY547-cGMP (fcGMP) previously shown to activate CNG channels with similar efficacy and affinity to cGMP (Biskup et al. 2007). To identify ZMW apertures containing single TAX-4 channels, we first estimated the number of ALFA–mSG–TAX-4 subunits in each aperture by photobleaching mSG under continual 488 nm excitation and counting the number of bleach steps in the mSG fluorescence time series at each ZMW location (**Fig. 2A**). Given that TAX-4 channels are tetramers, we excluded from analysis all locations with either zero or greater than four bleach steps indicative of either no channels or multiple channels, respectively. Next, we observed stepwise changes in fcGMP fluorescence time series at each ZMW location previously identified as having a single TAX-4 channel under continual 532 nm excitation (**Fig. 2B,C**). These steps in fluorescence intensity report on binding and unbinding of individual fcGMP molecules (Patel et al. 2021). Fluorescence trajectories for fcGMP were acquired for 5 minutes at 10 Hz.

**Figure 2.**
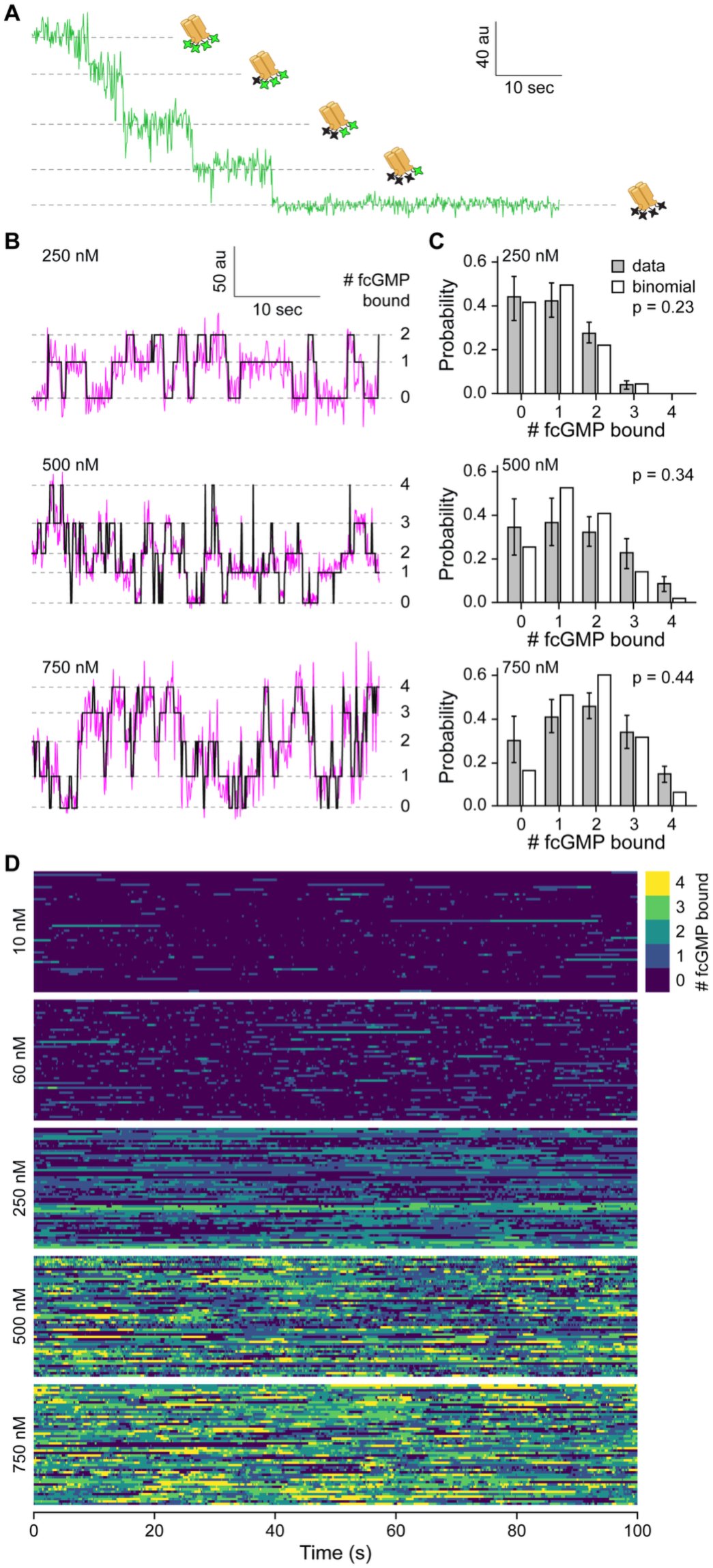
Stepwise binding of fcGMP to individual TAX-4 channels. (A) Fluorescence time series showing stepwise bleaching of mSG in a single ALFA–mSG–TAX-4 tetramer immobilized in a ZMW aperture. (B) Representative fluorescence time series (magenta) for three ZMWs showing stepwise binding and unbinding of fcGMP to TAX-4 at 250, 500, or 750 nM fcGMP. The idealized stepwise binding trajectory for occupation of 0-4 binding sites is overlaid (black). (**C)** Average concentration-dependent probability for each ligation state across all molecules and associated binomial fits for four independent and identical sites with bound probability *p* as indicated. Error bars are standard deviation across molecules **(D)** Graphical depiction of binding dynamics for ∼1.4 hours of binding at each concentration of fcGMP. Images are constructed from concatenated binding time series where each row represents single-channel binding dynamics.

Time series for fcGMP binding were recorded at 250, 500, and 750 nM fcGMP (**Fig. 2B,D**). The frequency of binding events increased with concentration as expected for a binding reaction. At 250 nM fcGMP, we mostly observed mono-and di-liganded states (i.e., one and two fluorescence steps). With increasing concentrations at 500 and 750 nM fcGMP, we observed tri-and quad-bound ligation states (i.e., third and fourth fluorescence steps) with increasing frequency. Thus, this dataset resolves all four sequential binding steps at TAX-4 channels. To evaluate whether the observed binding events reflect specific binding to TAX-4 channels as opposed to non-specific interactions with the ZMW surface, we compared binding trajectories in fcGMP alone with those in a mixture of fcGMP and an excess of non-fluorescent cGMP. The addition of cGMP resulted in a pronounced reduction in fcGMP binding events, consistent with competitive binding at TAX-4 channels (**Fig. S6**). This suggests that the observed binding events represent specific binding of fcGMP to TAX-4 CNBDs.

To quantify trajectories amongst discrete ligation states (i.e., number of bound ligands), we identified the set of discrete fluorescence levels in each fcGMP fluorescence time series using the divisive segmentation and clustering (DISC) algorithm (White et al. 2020) with automated per-molecule choice of information criterion (AutoDISC) (Bandyopadhyay and Goldschen-Ohm 2021). The idealization of each time series was manually inspected. In some cases, DISC was constrained to a visually identified number of levels—typically to account for a missed infrequently visited level. To better resolve dynamics, we further optimized each idealized trajectory using the Baum-Welch algorithm for a Markov model allowing all possible transitions amongst fixed fluorescence levels as identified by DISC (**Fig. 2B**).

For analysis, we combined data at lower fcGMP concentrations from our previous study (Patel et al. 2021) with data at 250, 500, and 750 nM fcGMP in this study (see Methods). The subsequent analysis below is based on this combined dataset, for which we provide a visualization of ∼1.4 hours of idealized binding dynamics at each of several concentrations of fcGMP (**Fig. 2D**). The per-molecule average bound probability increased with fcGMP concentration as expected for a binding reaction (**Fig. S7**).

### Cooperative binding across subunits

If binding at each subunit is identical and independent, the probability for 0-4 sites to be occupied by fcGMP should be binomially distributed. Binomial fits to the distribution of per molecule average observed probabilities in each ligation state indicate an increasing per-site occupation probability (*p*) with increasing fcGMP concentration from *p* = 0.23 at 250 nM to *p* = 0.44 at 750 nM, consistent with a binding reaction (**Fig. 2C**). These binomial bound probabilities are similar to those observed for fcAMP (an analogous fluorescent cAMP analog) binding to HCN channels at similar ligand concentrations (White et al. 2021). Although a binomial fit reasonably explains our observations at 250 nM fcGMP, our measurements deviate from a binomial distribution at higher ligand concentrations where we observe comparatively less time spent in mono-and di-liganded states and more time spent in tri-and quad-liganded states. This could reflect cooperativity amongst binding sites that promotes subsequent binding events.

Additional evidence supporting overall positively cooperative binding comes from first latencies to stepwise binding events in each distinct ligation state. Distributions of the latency with *i*-1 bound fcGMP ligands preceding binding of the *i^th^* ligand inform on the stepwise association dynamics (**Fig. S8**). For identical and independent binding sites, the effective binding rate is a product of the number of available sites. Thus, the latency to binding of the next ligand would be expected to increase with progressively fewer available sites during stepwise binding. In contrast, we observe decreasing latencies with increased occupation, suggesting that binding is positively cooperative (**Fig. S8**).

### Relating binding to channel activation

To assess the functionality of the ALFA–mSG–TAX-4–PS–TS fusion protein, we transiently expressed it in *Xenopus laevis* oocytes and recorded cGMP-and/or fcGMP-activation from excised inside-out patches (**Fig. 3A**). We chose oocytes for these experiments due to the relative ease of obtaining robust channel currents in excised patches (required to gain access to the cGMP binding domains) as compared to HEK-293T cells. Despite some differences in oocyte and HEK-293T cell membrane environments, concentration-response curves (CRCs) for cGMP-evoked current amplitudes were similar to those previously reported in mammalian cell membrane (Komatsu et al. 1999) (**Fig. 3B**, 𝐸𝐶_50_= 4.17 × 10^-7^ M, ℎ = 1.28, n = 5). Thus, we conclude that the fusion protein is activated by cGMP normally. In contrast, fcGMP CRCs were right-shifted by ∼2-fold with respect to cGMP CRCs, indicating a slightly lower affinity for fcGMP as compared to cGMP (**Fig. 3B**, 𝐸𝐶_50_ = 8.34 × 10^-7^ M, ℎ = 1.14, n = 4). Nonetheless, the concentration-dependence of fcGMP currents was essentially identical to that for the average bound probability from single-molecule measurements, suggesting that the binding dynamics observed in vesicles is relevant to that driving channel activation (**Fig. 3B**).

**Figure 3.**
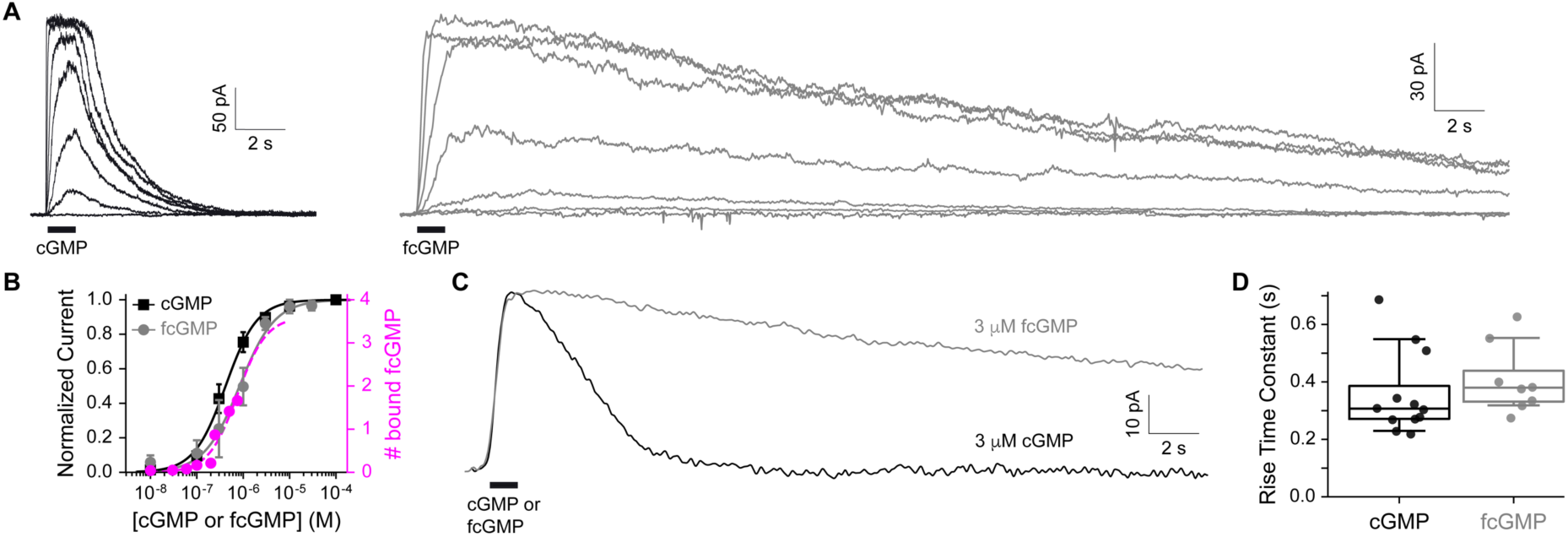
Relating channel activation to bound probability. **(A)** Families of current responses to 1 s pulses (black bars) of either cGMP or fcGMP applied to inside-out patches excised from oocytes expressing the ALFA–mSG–TAX-4–PS–TS fusion protein. Each family of responses is from a single patch for concentrations ranging from non-activating to maximally activating. **(B)** Normalized concentration-response curves for peak cGMP-or fcGMP-elicited currents (black squares, gray circles; mean ± sem across patches) and the average bound probability for fcGMP across single-molecule time series (magenta circles). Curves are fits to the means with the Hill equation (Eq. 1). **(C)** Comparison of current responses to 1 s pulses (black bars) of either cGMP or fcGMP applied to inside-out patches excised from HEK293T cells expressing the ALFA–mSG–TAX-4–PS–TS fusion protein. Both responses are from the same patch. **(D)** Summary of the time constant for the rise to peak current from HEK293T cell patches as shown in panel C. Data points are individual patches. Box plots are the median and quartiles.

However, current decay upon removal of fcGMP was much slower than for cGMP, persisting for tens of seconds following a 1 s pulse of fcGMP. To verify that this was not due to differential function in oocytes versus mammalian cells, we obtained recordings from inside-out patches of HEK-293T cells expressing the fusion construct in response to 1 s pulses of 3 μM cGMP or fcGMP, a concentration that is close to saturating. Current responses from HEK-293T cells recapitulated our observations from oocytes, exhibiting identical current activation kinetics for both cGMP and fcGMP, but significantly prolonged current decay with fcGMP (**Fig. 3C,D**). We conclude that fcGMP activates TAX-4 channels in much the same way as cGMP but unbinds much more slowly.

### Binding dynamics suggest positively cooperative binding followed by isomerization of individual binding domains

The slower unbinding kinetics of fcGMP as compared to cGMP imply that our observations of fcGMP binding dynamics will differ from those of cGMP. Nonetheless, the nearly identical current activation kinetics elicited by both fcGMP and cGMP suggest that the observed dynamics for stepwise activation by fcGMP are in some respects physiologically relevant to activation by cGMP. In fact, the longer bound lifetimes for fcGMP are expected to simplify resolution of binding dynamics which otherwise may struggle to resolve very short-lived events. Thus, we proceeded to analyze the binding dynamics by comparing the likelihood of several Markov models postulating different binding mechanisms.

We explored three types of binding mechanisms, each with either cooperative or identical and independent binding sites (**Fig. 4A**). The model subscript indicates whether binding was constrained to be identical and independent (X_i_) or allowed to be cooperative (X_c_). All models were optimized to maximize their likelihood for either the entire combined dataset at fcGMP concentrations from 10 to 750 nM (see methods), bootstrapped versions of this dataset, or each of three stratified folds of the dataset (**Table S1**). The models were ranked according to their relative Bayesian information criterion (ΔBIC) scores, which is an empirical measure that balances goodness of fit with model complexity (a lower ΔBIC score is better). Whether or not cooperative binding was allowed, models describing only sequential stepwise binding to four sites (B_i/c_) were ranked much worse than models that additionally allowed for an isomerization of the binding domain(s) distinct from the binding event itself (G_i/c_, S_i/c_) (**Fig. 4B**). This observation parallels our prior observations for early fcGMP binding steps at TAX-4 channels (Patel et al. 2021) and similar observations by ourselves and others for fcAMP binding to HCN channels or isolated binding domains (Goldschen-Ohm et al. 2016, 2017; White et al. 2021). We conclude that any reasonable model must account for both association events and isomerization of the ligand-bound domains.

**Figure 4.**
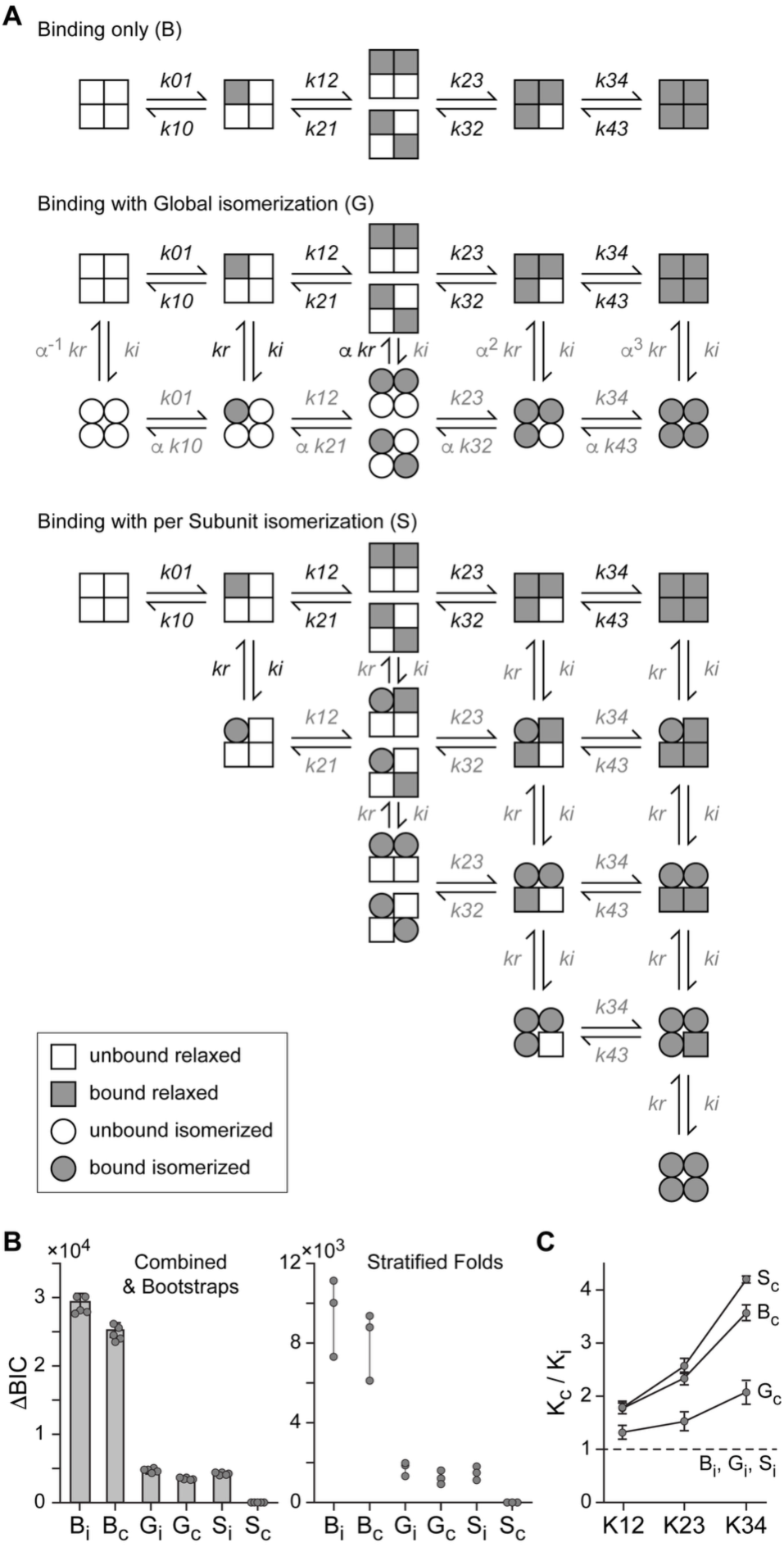
Binding dynamics suggest cooperative binding followed by an isomerization of individual sites. **(A)** Evaluated HMM model topologies include stepwise binding/unbinding only, binding with a global isomerization of all four binding domains, or binding followed by isomerization of individual binding domains. For each topology, binding/unbinding rates were either constrained as for independent and identical binding sites (*i* subscript) or allowed to be cooperative (*c* subscript) (see Table S1 for constraints and optimized parameters). **(B)** Models were ranked according to their relative Bayesian Information Criterion (ΔBIC) scores (lower is better). Similar patterns of ΔBIC scores were obtained from maximum likelihood fits to either a combined dataset including all tested fcGMP concentrations (left, bars), five bootstrap resampled datasets (left, circles), or each of three stratified folds of the dataset (right), suggesting that the fits were not overly skewed by only a part of the dataset. **(C)** The fold-change in the binding equilibrium constants (K = binding rate / unbinding rate) for cooperative models (K_c_) over that expected for independent and identical binding sites (K_i_) indicates a stepwise increase in positive binding cooperativity.

Amongst these models, we explored either a global isomerization of all four binding domains where binding promotes isomerization (G_i/c_) or per-subunit isomerization of individual binding domains following binding (S_i/c_). When binding was constrained to be identical and independent at all four sites, we could not distinguish between a global (G_i_) or per-subunit (S_i_) isomerization mechanism based on ΔBIC scores. In contrast, allowing for cooperative binding was of little benefit to the model with a global isomerization (G_c_), but resulted in our best ranked model when coupled with per-subunit isomerization of individual binding domains (S_c_). These results were robust across both bootstrapped datasets and three stratified folds of the combined dataset (**Fig. 4B**). When cooperative binding was allowed, we did not constrain the type of cooperativity (i.e., positive or negative) at each subsequent binding step. However, in all cases the models identified increasing positive cooperativity with sequential binding steps (**Fig. 4C**). We conclude that the most likely mechanism includes positively cooperative binding followed by isomerization of individual occupied binding domains.

## Discussion

Electrophysiological measures of ligand-gated ion channel activity report on the final ion conduction events, but do not directly reveal the sequence of ligand association events and conformational transitions that precede pore gating. Optical methods enable observation of electrically silent events, with fluorescently labeled ligands reporting on the beginning of the stimulus-response pathway in much the same way that ionic current reports on the end of this pathway. However, diffraction-limited microscopy limits typical optical approaches to low ligand concentrations, often below what is required for physiologically relevant activity. Here, we overcome this limitation using ZMWs to observe the sequential binding of fcGMP to each of four subunits in TAX-4 cyclic nucleotide-gated ion channels in cell-derived vesicles. Our approach combines methods that enable optical resolution of individual binding events at up to micromolar concentrations of fluorescent ligand with isolation of channels in native cell membranes without solubilization in detergent or polymers that can disrupt channel function.

The kinetics and cooperativity of ligand binding in CNG and related hyperpolarization-activated cyclic nucleotide-gated (HCN) channels have been investigated in different channel types such as vertebrate rod and olfactory CNG channels, the *C. elegans* TAX-4 channel, and several HCN channel isoforms (Goldschen-Ohm et al. 2016, 2017; White et al. 2021; Nache et al. 2005; Biskup et al. 2007; Nache et al. 2013; Idikuda et al. 2025; Chowdhury, Goldsmith, and Chanda 2026; Costa et al. 2026). These studies have yielded remarkably differing conclusions regarding the degree or even direction of coupling between sequential binding events and the connection between distinct ligation states and channel gating. Although fundamental differences between channel types may underlie some of these observations, even studies of the same channel type have reached apparently contradictory conclusions—possibly owing to different experimental approaches and their associated strengths and weaknesses. For example, studies which inferred the dynamics of the activation process from ensemble-averaged measures of binding or evoked channel current are challenged to uniquely resolve the asynchronous, molecule-specific binding sequences that underlie gating. Conversely, studies which have resolved the binding dynamics at single molecules were blind to the downstream gating of the channel and may be affected by non-native environments in the preparation such as solubilization in detergents. Ultimately, the reason for the disparate conclusions remains to be determined.

Our observations suggest that fcGMP binding to TAX-4 CNG channels in native cell membrane is a positively cooperative process, and that binding induces an isomerization of the occupied binding domain in individual subunits. Our conclusion of positive binding cooperativity parallels that for similar observations of single-molecule binding dynamics at HCN channels in lipid bilayers (Idikuda et al. 2025). In contrast, no binding cooperativity was observed when HCN was solubilized in detergent (White et al. 2021), highlighting the value of a native membrane preparation. Although our best-ranked model suggests that each binding domain isomerizes individually following ligand binding, this isomerization is too slow to reflect gating of the channel pore. Open state lifetimes from single channel recordings are on the millisecond time scale as opposed to the second time scale for isomerization events observed here (X. Zheng et al. 2022). We interpret this isomerization as a pre-active intermediate along the activation pathway. This asynchronous isomerization of individual binding domains could culminate in a global conformational change such as a concerted movement of the C-linkers which may be more directly associated with pore gating or even periods of high activity such as during bursts of channel openings, but which we do not directly observe. Although such a model seems plausible, the additional parameters involved and the inability of our binding-centric observations to directly constrain them makes exploration of such models with our dataset largely infeasible.

Our analysis is complicated primarily by two factors. The first is signal-to-noise, especially when multiple ligands are bound simultaneously due to additive noise from multiple fluorophores, which is likely to be a limiting factor in our analysis. The second is the slow unbinding of fcGMP as compared to cGMP (**Fig. 3**). Macroscopic simulations of the average bound probability in response to a 1 s pulse of fcGMP using our preferred model (S_c_) are consistent in their prediction of slower unbinding than expected for cGMP based on our observed current decay kinetics. However, the predicted fcGMP unbinding time course is still faster than the observed current decay following activation by fcGMP. Thus, it is likely that our binding data are contaminated by fluorophore bleaching due to the prolonged lifetime of bound events. Despite these complications, activation kinetics are identical for both cGMP and fcGMP (**Fig. 3C,D**), and our tested models strongly support a mechanism involving positive binding cooperativity and an isomerization of the binding domains independent from the association events themselves is likely (**Fig. 4A,B**). Furthermore, even the simplest binding-only model (B_c_) identified similar stepwise binding cooperativity to our preferred model including isomerization of occupied sites (S_c_) (**Fig. 4C**). This suggests that even simplistic analysis of these data can still reveal fundamental aspects of the activation mechanism.

Ultimately, simultaneous measures of both ligand binding and channel gating dynamics at single-molecule resolution may be needed to provide a clearer picture of the full activation energy landscape. Alternatively, coupling binding measures with other perturbations having known effects on gating or orthogonal optical readouts of conformational changes could enhance mechanistic insight. Either way, the approach illustrated here provides a broadly applicable tool to dissect the energetic landscape of ligand-binding and associated early conformational changes at macromolecules in native cell membranes.

## Methods

### Construct

DNA for the TAX-4 construct used in this study was synthesized and subcloned in the pUNIV vector (Venkatachalan et al. 2007) by GenScript. The has an ALFA tag followed by the fluorescent protein mStayGold (mSG) fused to the N-terminus, and a PreScission (PS) cleavage site followed by a Twin-Strep® (TS) tag fused to the C-terminus (ALFA–mSG–TAX-4–PS–TS; Fig. 1). Complementary RNA (cRNA) for each construct was generated (mMessage mMachine T7, Ambion) and quantified (Qubit, ThermoFisher Scientific) prior to injection in *Xenopus laevis* oocytes.

### Preparation of cell-derived vesicles

HEK-293T cells (ATCC) were cultured at 37°C and 5% CO_2_ (Eppendorf). Cells were plated in 60 mm dishes and transfected with 1.5 μg of ALFA–mSG–TAX-4–PS–TS and 3 μg of PEI-MAX per dish. After 48 hours, cells from 12 dishes were combined and vesicles were prepared as previously described (Moonschi et al. 2018; Fox-Loe, Moonschi, and Richards 2017; Patel et al. 2021). Briefly, cells were disrupted by nitrogen cavitation at 600 psi for 20 minutes (Parr Instrument Company #4639) while suspended in 3 ml of a hypotonic buffer (in mM: 10 Tris-HCl, 10 NaCl, 1.5 MgCl_2_, 0.2 CaCl_2_, pH 7.4) including one Pierce protease inhibitor tablet (ThermoScientific #A32955) per 10 ml of buffer. The membrane fragments naturally form vesicles due to the hydrophobicity of the lipids. Plasma membrane vesicles were separated from other organelle membranes such as endoplasmic reticulum using gradient ultracentrifugation in a swing bucket rotor (Beckman SW41Ti rotor, 13.2 ml tubes #344059) at 112,000×g for 90 minutes at 4 °C (Beckman Optima XPN 80). The gradient was prepared with 30, 20, and 10% solutions of OptiPrep (Sigma-Aldrich #D1556-250ML) diluted with sucrose buffer (in mM: 250 sucrose, 10 HEPES, pH 7.4). The isolated plasma membrane fraction was pelleted by ultracentrifugation at 100,000×g for 1 hour at 4 °C using a fixed angle rotor (Beckman 70Ti rotor, 6.5 ml tubes #344088) and then resuspended in 300 μl of wash buffer consisting of phosphate-buffered saline (PBS, pH 7.4) supplemented with 1 mM EDTA, 0.5% BSA, and 10% glycerol as per the magnetic bead protocol described below.

Vesicles containing ALFA–mSG–TAX-4–PS–TS oriented with the TS tag exposed to the bath were purified using Strep-Tactin® Magnetic Microbeads (IBA Lifesciences #6-5510-050) according to the manufacturer’s protocol. Briefly, the microbeads were prepared as described by the manufacturer. Next, 300 μl vesicle sample was incubated with 350 μl of microbeads for 20 minutes at 4°C in a 1.5 ml tube under gentle constant agitation on a nutator. The beads were transferred to a 15 ml tube, and unbound vesicles were removed by washing with 8.5 ml of wash buffer three times (25.5 ml total wash) using a magnet to separate the supernatant from the beads after each wash. The beads were transferred back to a 1.5 ml tube, and bound vesicles were separated from the beads by incubation with PreScission enzyme (1.4 units/ul for ∼14 ul of beads) in 1 ml cleavage buffer (in mM: 50 TrisHCl, 150 NaCl, 1 EGTA, 1 DDT, 10% glycerol) for 16 hours at 4°C under constant agitation on a nutator. This cleaves the fusion protein at the PS motif. Eluted vesicles containing ALFA–mSG–TAX-4 were separated from the beads using a magnet. We repeated the magnetic separation three times to ensure no beads were collected.

To constrain the vesicle size distribution for loading into ZMWs, vesicles were extruded by passing between two 1 ml syringes through a nanopore (Avanti Polar Lipids). To ease manual extrusion, the eluted vesicles were diluted in PBS to a total volume of 1 ml. Vesicles were then extruded through a 50 nm pore (Whatman #800308) for 40 passes (i.e., 20 times through and back). Finally, the vesicles were extruded through a 30 nm pore (Whatman #800307) for 2-4 passes before loading onto ZMW arrays.

### Imaging fcGMP binding at TAX-4 channels in ZMWs

Zero-mode waveguide (ZMW) arrays (Pacific Biosciences) whose bottom glass coverslip is passivated with a biotin-doped polyethylene glycol (PEG) layer were sequentially incubated in 10 mg/mL bovine serum albumin (BSA), followed by 0.05 mg/mL streptavidin (Fisher Scientific #434302), and finally 1 µg/ml biotinylated anti-ALFA nanobody (NanoTag Biotechnologies #N1505). This procedure functionalizes the bottom surface with anti-ALFA nanobodies via biotin-streptavidin interactions. All incubation solutions were prepared in phosphate-buffered saline (PBS; pH 7.4). Following each incubation step (10–15 minutes at room temperature), the ZMW chamber was washed five times with PBS supplemented with 2 mg/ml BSA to remove unbound material. Extruded vesicles containing ALFA–mSG–TAX-4 were deposited onto the functionalized ZMW arrays by incubating for 30 minutes, after which unbound vesicles were removed by washing 5-10 times as described above. Finally, the buffer was exchanged for imaging buffer consisting of PBS supplemented with 1 mM Trolox, 2.5 mM protocatechuic acid (PCA), and 250 nM protocatechuate 3,4-dioxygenase (PCD; from *Pseudomonas* sp.) to create an oxygen-scavenging environment and help protect against fluorophore bleaching. The imaging buffer also contained various concentrations of 8-(2-[DY-547]-aminoethylthio) guanosine-3’,5’-cyclic monophosphate (fcGMP; BioLog).

The chamber of ZMW arrays was imaged on a custom inverted microscope (Mad City Labs RM21) with 60× magnification (Olympus APON 60X TIRF objective). Fluorescence from mSG or fcGMP in ZMW apertures was recorded on a sCMOS camera (Teledyne Photometrics Prime BSI Express) at a frame rate of 10 Hz upon laser excitation through the objective using 488 nm or 532 nm epi illumination (Coherent OBIS). Micro Manager was used for image acquisition. Fluorescence emission from mSG or fcGMP was bandpass filtered at 500-550 nm or 542-950 nm, respectively (Chroma). A glass coverslip was placed over the top of the chamber to effectively minimize evaporation of the ∼70 µl chamber volume during recordings.

Fluorescence time series for individual ZMWs were extracted from image time series as the mean pixel intensity per frame for all pixels within an eight-pixel diameter circular mask centered on the ZMW aperture. Mechanical drift during each recording was subpixel due to the stability of the RM21 stage and thus was ignored. A small offset of up to a few pixels was sometimes observed between mSG and fcGMP image sets due to mechanical perturbation from manual swapping of filters on the optical table. This offset was corrected by registering brightfield images of the ZMW arrays for mSG and fcGMP recordings using an affine transform. Initially, all mSG fluorophores were bleached under 488 nm excitation until no obvious mSG spots remained (typically 1-2 minutes). Projections and registration were done with a custom plugin in the Python image analysis program napari. Bleach steps for mSG at each ZMW aperture were manually evaluated to estimate the number of ALFA–mSG–TAX-4 subunits in each spot. Spots with more than four mSG bleach steps indicative of multiple channels were excluded from analysis. Next, fcGMP fluorescence reporting on stepwise binding to TAX-4 channels was recorded under 532 nm excitation for at least five minutes. All recordings were obtained at a frame rate of 10 Hz. An initial time-dependent decay in fcGMP fluorescence which we attribute to bleaching of fluorescence contaminates in the sample chamber was observed across the entire image plane including locations outside of ZMW apertures. For each individual ZMW, we corrected for this decay by subtracting a spline fit to the background fluorescence time series at a location just outside of the ZMW. At some locations exhibiting mSG fluorescence, we did not observe clear fcGMP binding events. Although the reason for this is unknown, we hypothesize that surface interactions between the vesicle or channel protein may have rendered these channels either inaccessible or functionally impaired. These locations were excluded from further analysis.

### Idealization of individual binding events

Background-subtracted fcGMP fluorescence time series were denoised with AutoDISC (Bandyopadhyay and Goldschen-Ohm 2021) to identify fluorescence levels associated with stepwise binding to TAX-4 channels. Prior to denoising, infrequent artifacts attributed to adsorption of fcGMP to the bottom surface of a ZMW aperture were manually masked out. These events typically have much higher fluorescence than binding steps and thus are usually readily identifiable as described previously (Patel et al. 2021). All the denoised traces were visually inspected. In some cases, this approach either failed to resolve a visually apparent level—typically for levels visited only infrequently—or identified an additional level visually classified as noise. In such cases, we constrained DISC to the visually identified number of levels. Any trajectories exhibiting more than five levels (i.e., more than four binding steps) were excluded from further analysis, as were trajectories with low signal-to-noise which challenged reliable state assignment. Differences in signal-to-noise could reflect variation in the physical geometry of individual ZMWs, their passivation, and non-uniform excitation across the field of view from the gaussian laser beam profile. Finally, the dynamics of each idealized trajectory was optimized using the Baum-Welch algorithm for a Markov model allowing all possible transitions amongst fixed levels as identified by DISC.

### HMM models

For a more comprehensive dataset, we combined single-molecule binding trajectories at 250, 500, and 750 nM fcGMP as described here with trajectories at 10, 30, 60, 100, and 200 nM fcGMP as we reported previously (Patel et al. 2021). To avoid bias towards dynamics at lower concentrations where we have more data, we selected a subset of the lower concentration data such that the total duration of dynamics at each concentration is similar. In total, we recorded ∼29 hours of single-molecule binding (∼3.6 hours at each concentration). Hidden Markov models were optimized for the entire combined dataset of idealized binding dynamics using QuB (Nicolai and Sachs 2013).

### Electrophysiology

Defolliculated *Xenopus laevis* oocytes were obtained from Ecocyte. Oocytes were injected with 25 ng of cRNA encoding the ALFA–mSG–TAX-4–PS–TS fusion construct and incubated in a sterile incubation solution (88 mM NaCl, 1mM KCl, 2.4 mM NaHCO_3_, 19 mM HEPES, 0.82 mM MgSO_4_, 0.33 mM Ca(NO_3_)_2_, 0.91 mM CaCl_2_, 10,000 units/L penicillin, 50 mg/L gentamicin, 90 mg/L theophylline, and 220 mg/L sodium pyruvate, pH 7.5) at 16 °C for 1-2 days. Immediately prior to patch-clamp recording, the oocyte vitelline membrane was removed using forceps after incubating it in stripping solution (200 mM potassium aspartate, 20 mM KCI, 1 mM MgCI, 10 mM EGTA, 10 mM HEPES, pH 7.4) for 5-10 min.

HEK-293T cells (ATCC, CRL-3216) were cultured in MEM (Gibco, 11095-080) supplemented with 10% fetal bovine serum (Gibco, A5256701), 1% non-essential amino acids (Gibco, 11140-050), 1% sodium pyruvate (Gibco, 11360-070) and 1% penicillin/streptomycin (Gibco, 151140-122) at 37 °C and 5% CO_2_. For electrophysiological experiments, cells were plated on glass coverslips (Assistent, 92100100030) and transfected at ∼60–80% confluency using FuGENE 4k (Promega, E951A) with a total of 1 μg of DNA encoding the ALFA–mSG–TAX-4–PS–TS fusion construct per 35 mm dish of cells. Recordings were performed 24-48 hours post-transfection.

Excised inside-out patch recordings from either oocytes or HEK-293T cells were obtained at room temperature using an Axopatch 200A amplifier (Axon Instruments), ITC-16 digitizer (HEKA), and WinWCP acquisition software (Strathclyde). Cells were voltage-clamped at - 60 mV and recorded currents were lowpass filtered at 1 kHz. Patch pipettes (3-6 MΩ) were pulled (Sutter, P-97-4511) from borosilicate glass (Sutter, BF150-86-10). Recordings in oocytes were obtained in symmetrical intracellular and extracellular buffer solutions. Buffer solution for oocytes contained (in mM): 67 KCl, 30 NaCl, 10 HEPES, 10 EGTA, and 1 EDTA (pH 7.2 with KOH). The internal buffer solution for HEK-293T cells contained (in mM): 10 CsCl, 10 HEPES, 10 EGTA, 2 MgATP, and 110 CsF (pH 7.3 with KOH, 300 mOsm adjusted with sucrose as needed). The external buffer solution contained (in mM): 130 NaCl, 4 KCl, 10 CaCl₂, 1 MgCl₂, 10 HEPES, and 5 glucose (pH 7.3 with NaOH, 300 mOsm adjusted with sucrose as needed). cGMP and fcGMP were dissolved in the buffer solution and applied to inside-out patches via a multi-barrel perfusion system (Warner Instruments, SF-77B). Defined pulses of ligands were applied using a computer controlled piezoelectric stepper to rapidly switch perfusion barrels between buffer and ligand. Solution flow was controlled with a microfluidic pump (Elveflow, OB1-MK4). To obtain concentration-response curves for individual cells, the concentration of cGMP or fcGMP in one of the perfusion barrels was changed using a microfluidic switch (Elveflow, MUX-Flow).

Concentration-response curves (CRCs) were fit with the Hill equation:

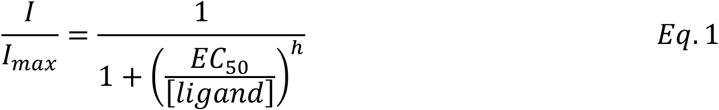

where 𝐼 is the magnitude of the ligand-elicited current, [𝑙𝑖𝑔𝑎𝑛𝑑] is cAMP or fcAMP concentration, 𝐸𝐶_50_is the concentration eliciting a half-maximal response, and ℎ is the Hill slope.

## Acknowledgements

This research was supported by NIH grant R01GM148591 to M.P.G-O. and additional support was provided by the Albany, Gottesman, and Wilder Foundations to A.C.B.

## Author Contributions

M.P.G.-O. conceived, designed, and supervised this work. T.H. carried out all purification, imaging, and analysis for single-molecule binding experiments. C.M.B. designed the construct and purification strategy, and transcribed RNA and injected oocytes. P.A.P.P. and S.R. generated preliminary constructs for this work. T.H. and Z.A. performed sample preparation.

D.W. carried out, and M.P.G.-O. and A.C.B. supervised, all electrophysiology recordings. T.H., C.M.B., D.W., and M.P.G.-O. wrote the manuscript.

---

**Figure S1.**
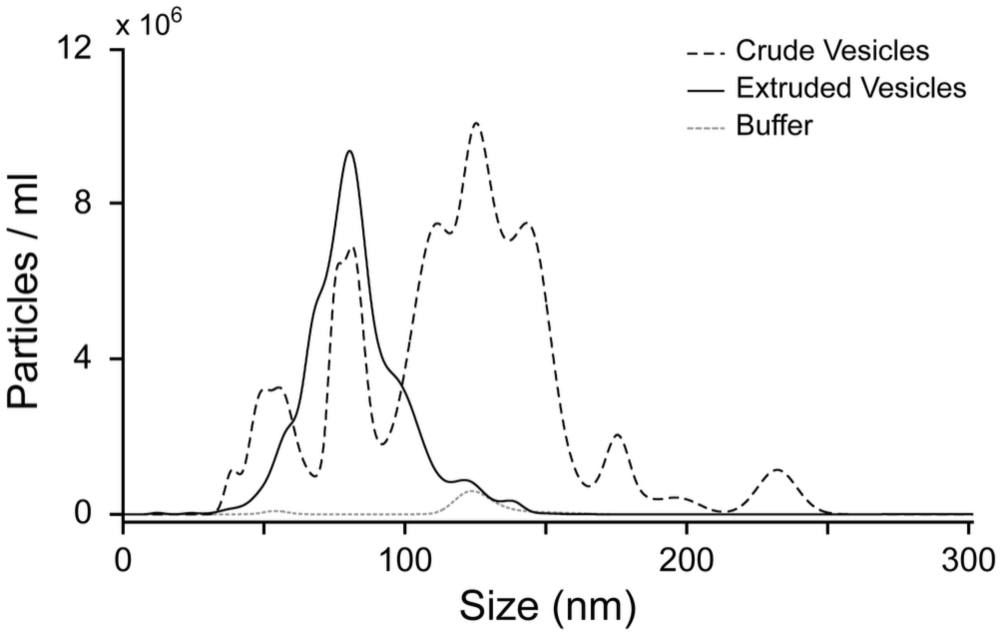
Extruded vesicle size distribution. Size distributions for non-extruded (crude, dashed line) or extruded (solid line) cell-derived vesicles. Control is for PBS buffer alone which had minimal particulate contaminants. Vesicle samples contained the TAX-4 fusion protein as described in the main text. Size distributions were obtained by nanoparticle tracking analysis (NTA) using a NanoSight NS500 (NanoSight, Amesbury, United Kingdom) equipped with a 488 nm laser.

**Figure S2.**
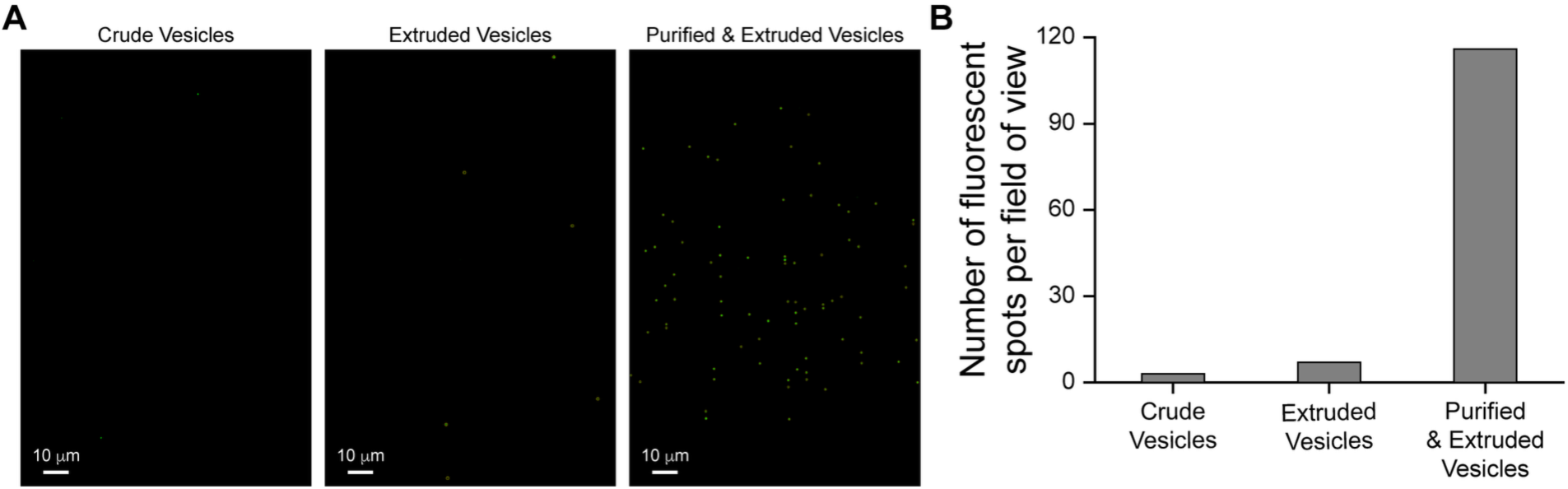
Loading TAX-4 containing cell-derived vesicles into ZMW arrays. **(A)** Fluorescence images of 150 nm ZMW arrays after loading with either crude or extruded vesicles containing the fusion protein eGFP–TAX-4 (left, middle), or purified and extruded inside-out cell-derived vesicles containing the fusion protein ALFA–mSG–TAX-4. Fluorescence is from mSG (right). **(B)** Each fluorescent spot is indicative of a ZMW well containing a vesicle with a TAX-4 fusion protein. Relatively efficient loading was only observed for samples that were both purified and extruded.

**Figure S3.**
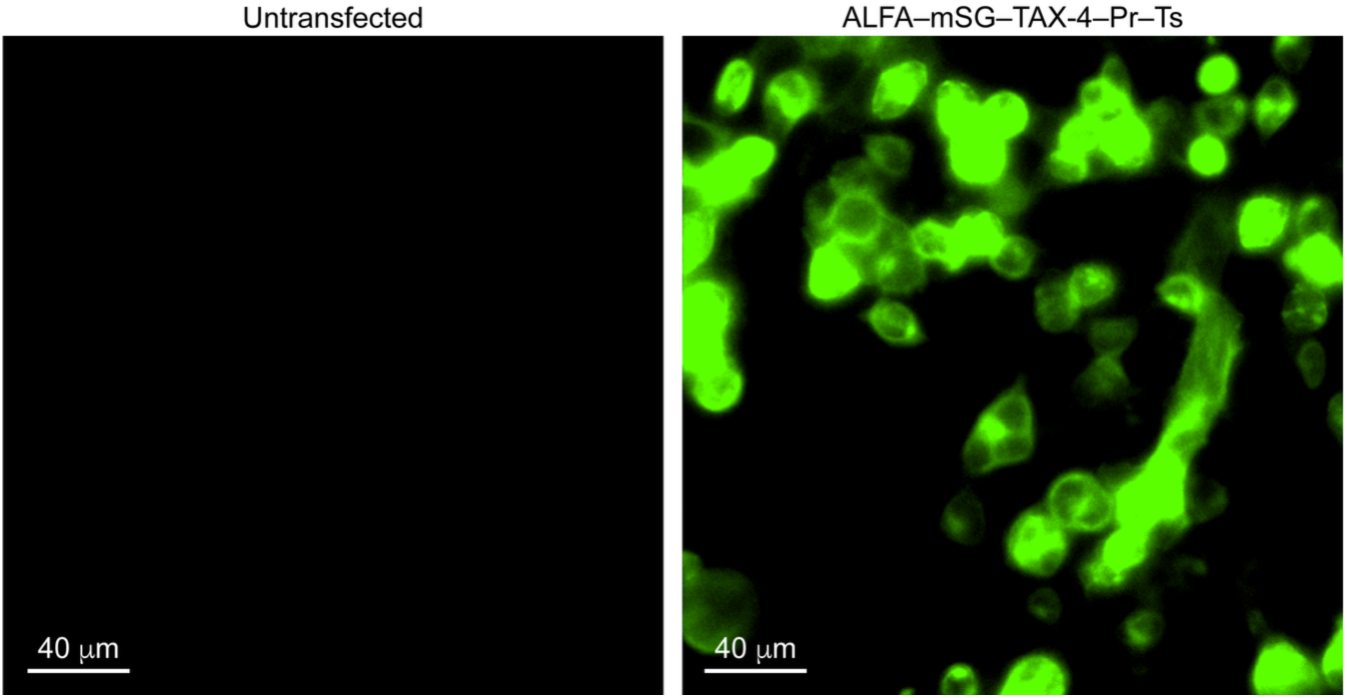
Expression of the fusion protein ALFA–mSG–TAX-4–PS–TS in HEK293T cells. Fluorescence emission from mStayGold (mSG) excited at 488 nm in HEK-293T cells either untransfected (left) or transfected with the fusion protein ALFA–mSG–TAX-4–PS–TS (right).

**Figure S4.**
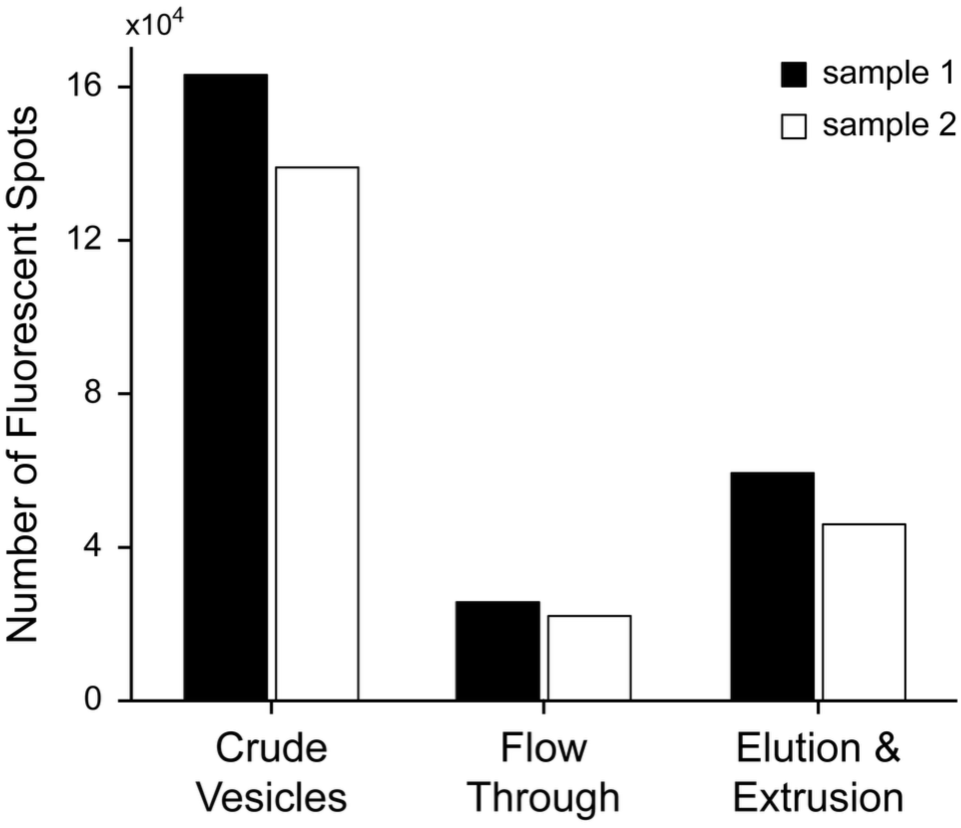
Purification of ALFA–mSG–TAX-4 in cell-derived vesicles. Fluorescence from mSG was used to quantify the amount of vesicles containing ALFA–mSG–TAX-4 during the purification process. We started with a 320 µl suspension of crude vesicles from the initial vesicle preparation which is comprised of vesicles both with and without the TAX-4 fusion protein in both inside-out and outside-out configurations. A 10 µl aliquot of the crude suspension was placed on a glass coverslip and imaged with TIRF illumination at 488 nm. The total number of fluorescent spots (e.g., vesicles) in a single field of view (F) was recorded and scaled by 32 to estimate the same measure for the entire sample. The remaining 310 µl was taken forward for purification and incubated with Step-Tactin coated magnetic beads. The beads were then washed with buffer to remove unbound vesicles and the flow through collected. A 50 µl aliquot of the flow through (300 µl total) was imaged as described for the crude vesicles and F was scaled by 6 to estimate the same measure for the entire flow through. Finally, the vesicles remaining bound to the beads were eluted by enzymatic cleavage at the PreScission site and the elute was collected and extruded in a final volume of 750 µl. A 50 µl aliquot of the extruded sample was imaged as described for the crude vesicles and F was scaled by 15 to estimate the same measure for the entire eluted and extruded sample. Scaled values of F at each step are shown for two separate samples on different days. Most of the TAX-4 containing vesicles remained bound to the beads, with only a small fraction ending up in the flow through. Roughly half of the bead-associated vesicles were eluted in the purified sample. Purification efficiency was similar for both samples.

**Figure S5.**
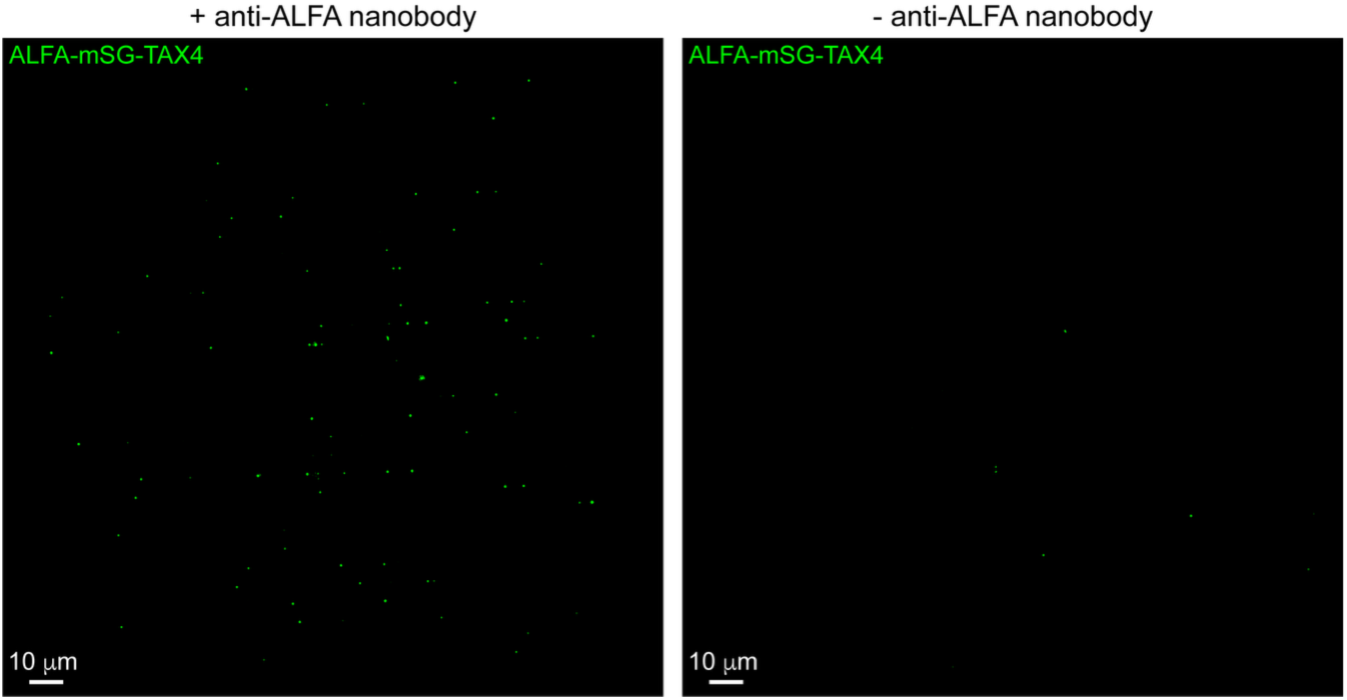
Specific immobilization of ALFA–mSG–TAX-4 containing vesicles. Fluorescence images of a ZMW array with (left) or without (right) anti-ALFA nanobodies on the bottom of the nano-wells after loading with cell-derived vesicles containing ALFA–mSG–TAX-4. Fluorescence from mSG is rarely observed in the absence of the nanobody, suggesting that mSG spots largely reflect specific immobilization via anti-ALFA nanobodies as opposed to non-specific adsorption of vesicles.

**Figure S6.**
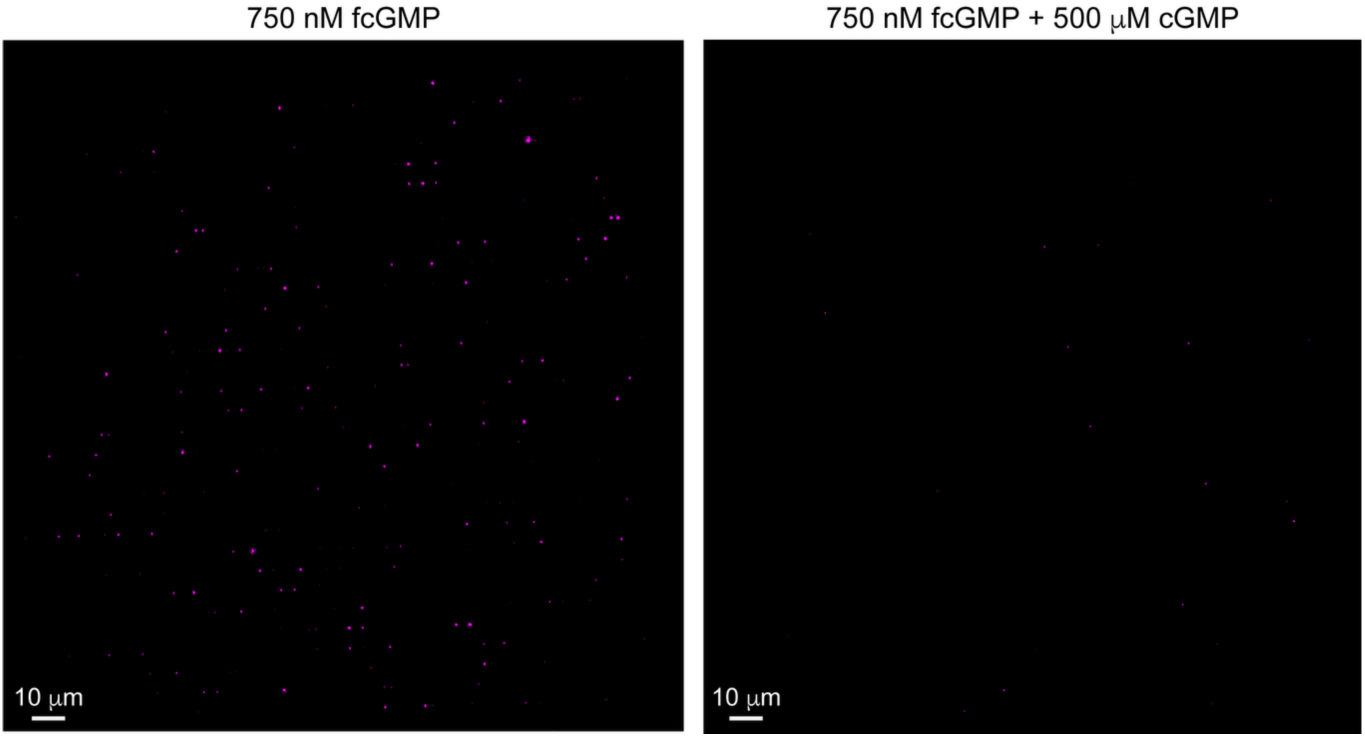
Specific binding of fcGMP to TAX-4 channels in cell-derived vesicles within ZMWs. Time-averaged fluorescence images for fcGMP binding. Both images are for the same field of view. Robust binding in 750 nm fcGMP (left) is largely abolished by competition with an excess of non-fluorescent cGMP (right).

**Figure S7.**
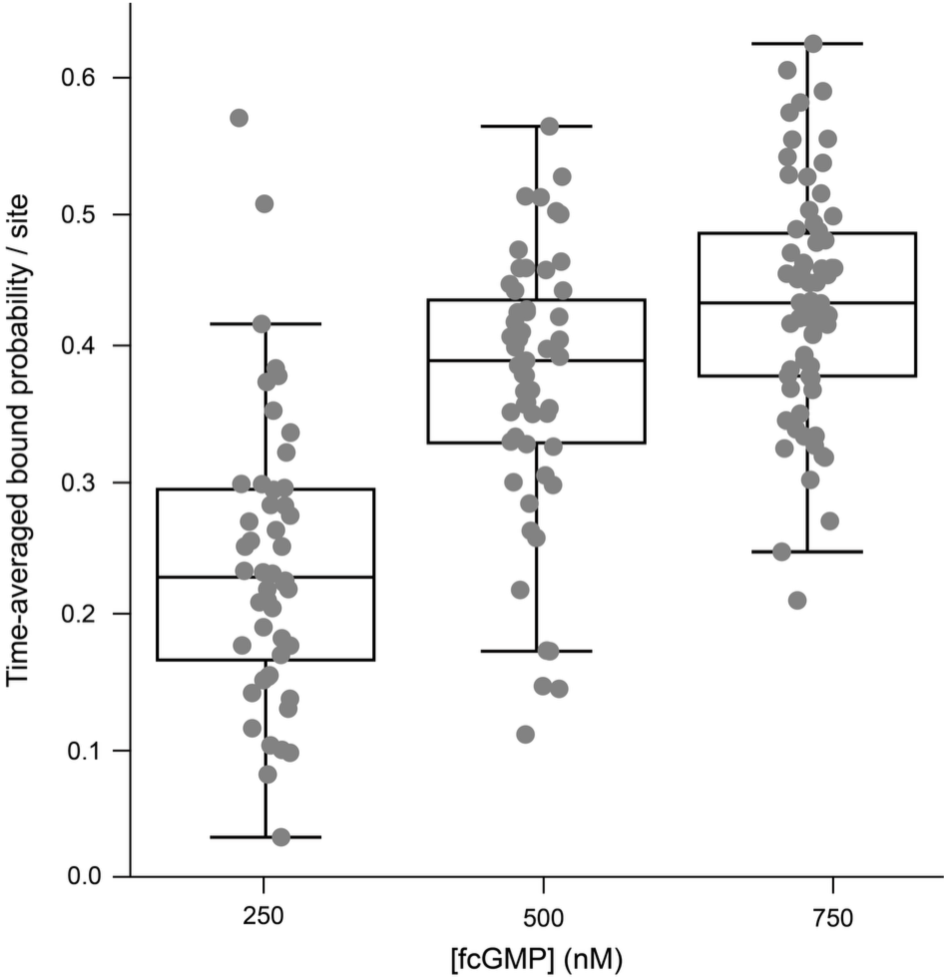
Probability of binding site occupation as a function of fcGMP concentration Each solid circle is the time-averaged bound probability per site for an individual TAX-4 channel. Box plots indicate the median and quartiles. The binding probability increases with increasing fcGMP ligand concentration, demonstrating a concentration-dependent interaction between TAX-4 containing nanovesicles and the fcGMP ligand.

**Figure S8.**
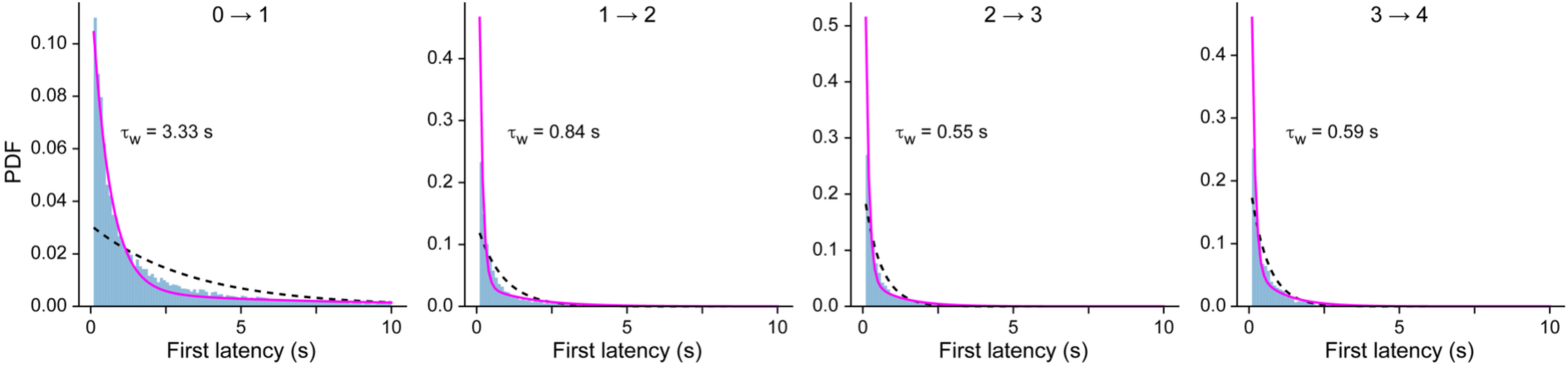
Step-wise binding first latencies. Distributions of first latencies to fcGMP binding for stepwise binding at four binding sites in TAX-4 channels. Mono-exponential (black dashed line) and bi-exponential (solid magenta line) maximum likelihood fits are shown, and the weighted time (τ_w_) constant from bi-exponential fits is shown.

**Table S1.** Model parameters. Parameters for kinetic models as shown in Fig. 4 of the main text. Units are μM^-1^s^-1^ for binding rates and s^-1^ for unbinding and isomerization rates, and 𝛼 is unitless. Parameters are from a global fit to a combined dataset including data at 10, 30, 60, 100, 200, 250, 500, and 750 nM fcGMP. Data at 250-750 nM fcGMP are from this work, and data from 10-200 nM fcGMP are from our previous publication (Patel et al. 2021). See methods in the main text. Error is the standard deviation for separate fits to three unique stratified folds of the combined dataset with approximatley equal representation at each concentration. Constraints are indicated by shaded cells. Degrees of freedom (*dof*) is the number of free parameters in each model and ΔBIC are the models’ relative Bayesian Information Criterion scores (lower is better).

| | Binding ($\mu\text{M}^{-1}\text{s}^{-1}$) | | | | Unbinding ($\text{s}^{-1}$) | | | |
| --- | --- | --- | --- | --- | --- | --- | --- | --- |
| Model | $k_{0\rightarrow1}$ | $k_{1\rightarrow2}$ | $k_{2\rightarrow3}$ | $k_{3\rightarrow4}$ | $k_{4\rightarrow3}$ | $k_{3\rightarrow2}$ | $k_{2\rightarrow1}$ | $k_{1\rightarrow0}$ |
| B<sub>i</sub> | $4k_{3\rightarrow4}$ | $3k_{3\rightarrow4}$ | $2k_{3\rightarrow4}$ | $0.7 \pm 0.2$ | $4k_{1\rightarrow0}$ | $3k_{1\rightarrow0}$ | $2k_{1\rightarrow0}$ | $0.7 \pm 0.2$ |
| B<sub>c</sub> | $2.3 \pm 0.4$ | $2.2 \pm 0.6$ | $1.7 \pm 0.6$ | $1.0 \pm 0.4$ | $1.7 \pm 0.6$ | $1.7 \pm 0.6$ | $1.2 \pm 0.3$ | $0.9 \pm 0.2$ |
| G<sub>i</sub> | $4k_{3\rightarrow4}$ | $3k_{3\rightarrow4}$ | $2k_{3\rightarrow4}$ | $0.7 \pm 0.2$ | $4k_{1\rightarrow0}$ | $3k_{1\rightarrow0}$ | $2k_{1\rightarrow0}$ | $2.0 \pm 0.4$ |
| G<sub>c</sub> | $2.4 \pm 0.5$ | $2.2 \pm 0.7$ | $1.7 \pm 0.6$ | $1.0 \pm 0.4$ | $6.4 \pm 1.1$ | $5.7 \pm 1.3$ | $3.8 \pm 0.7$ | $2.0 \pm 0.4$ |
| S<sub>i</sub> | $4k_{3\rightarrow4}$ | $3k_{3\rightarrow4}$ | $2k_{3\rightarrow4}$ | $0.7 \pm 0.2$ | $4k_{1\rightarrow0}$ | $3k_{1\rightarrow0}$ | $2k_{1\rightarrow0}$ | $1.9 \pm 0.4$ |
| S<sub>c</sub> | $2.5 \pm 0.5$ | $2.3 \pm 0.7$ | $1.8 \pm 0.7$ | $1.0 \pm 0.4$ | $3.9 \pm 1.4$ | $4.2 \pm 1.3$ | $3.5 \pm 0.9$ | $2.6 \pm 0.5$ |

**Table S1. Model parameters.**
| | Isomerization ($\text{s}^{-1}$) | | | | |
| --- | --- | --- | --- | --- | --- |
| Model | $k_i$ | $k_r$ | $\alpha$ | $dof$ | $\Delta\text{BIC}$ |
| B<sub>i</sub> | - | - | - | 2 | 29,212 |
| B<sub>c</sub> | - | - | - | 8 | 25,115 |
| G<sub>i</sub> | $0.1 \pm 0.02$ | $0.3 \pm 0.01$ | $0.16 \pm 0.01$ | 5 | 4,339 |
| G<sub>c</sub> | $0.1 \pm 0.02$ | $0.2 \pm 0.02$ | $0.17 \pm 0.02$ | 11 | 3,161 |
| S<sub>i</sub> | $0.4 \pm 0.03$ | $0.3 \pm 0.03$ | - | 4 | 3,983 |
| S<sub>c</sub> | $0.4 \pm 0.05$ | $0.2 \pm 0.03$ | - | 10 | 0 |

